# A response to Zhang et al. (2018), “Can Mouse-tracking Reveal Attribute Processing Speeds in Dietary Self-control? Commentary on Sullivan et al. (2015) and Lim et al. (2018)”

**DOI:** 10.1101/572974

**Authors:** Nicolette Sullivan, Cendri Hutcherson, Alison Harris, Antonio Rangel

**Affiliations:** Department of Psychology and Neuroscience, Duke University; Department of Psychology, University of Toronto; Department of Psychology, Claremont McKenna College; Humanities and Social Sciences, California Institute of Technology

## Introduction

In Sullivan et al. (2015), mouse-tracking was used in a food choice paradigm to test two related hypotheses: 1) that there are differences in the relative speed with which the decision-making circuitry computes and weights the value of attributes like health and taste; and, 2) that individual differences in these relative speeds are associated with individual differences in the ability to make healthy dietary choices. Consistent with these hypotheses, a regression analysis of the mouse-tracking paths found that health became significantly predictive of the mouse’s angle of movement ~195 ms later than taste, on average. Moreover, individual dietary self-control varied with estimates of the individual differences in the relative speed with which taste (tTaste) and health (tHealth) entered the decision-making process. Similar results have been found in other studies, including Lim et al. (2018) and the new data presented in Zhang et al. (2018).

### Summary of Zhang et al. (2018)’s Critique

Zhang et al. (2018) argue that the conclusions of the previous work are flawed due to problems with the methods used in the previous literature. Their criticism has three main components:

*C1. Choice-confound*. The authors argue that the results of the previous papers are flawed because the key regressions of the mouse-tracking paths need to include the actual choice direction (left or right) as a regressor to avoid confounds. For example, in p.5 they state that “the methodological problem becomes clear if one realizes that the correlations between the attributes and trajectory angle are confounded by actual choice”. To address this concern, they run the path regressions with the choice control and find the previous results for tHealth and tTaste disappear, which they take as evidence for a problem with the original methodology.
*C2. Curvature-argument*. Using the data from a related dietary choice experiment from their labs, the authors find a correlation between the estimates of tTaste obtained using our method and a measure of path curvature (maximum deviation, MD) in a “control or filler” task. They state that (p. 16) “if the processing speeds in Sullivan et al. (2015)’s method were derived from the genuine influences of attributes, they should not correlate with personal mean MD in the filer trials”. They use this to conclude that the findings in the previous papers are due to a motor-movement confound.
*C3. Non-robust statistical methodology*. An important technical challenge in the methodology proposed in Sullivan et al. (2015) is to minimize potential biases in the estimates of tTaste and tHealth driven by the fact that taste is weighted more heavily than health in the overall decision-making process of the typical individual. To address this concern, Sullivan et al. (2015) employed two different methods to estimate tTaste and tHealth. The authors here find that the results in Sullivan et al. (2015) replicate in their data using one method but not another, and they take this as further evidence for a problem with the original methodology.

Based on these three points, they conclude that their “re-analyses and additional data demonstrate that the seemingly strong evidence from their correlational analyses is severely inflated by a motor-control factor operating as a confounding variable”.

### Critical Evaluation of Zhang et al. (2018)

We agree with the authors that the self-control mechanisms hypothesized in Sullivan et al. (2015), if correct, are potentially important for understanding and improving self-control in several domains. This, together with the fact that other groups are building on this work, makes it important to understand any potential methodological shortcomings.

We also agree with the authors that the methodology proposed in Sullivan et al. (2015) can be improved. More work needs to be done to pin down the computational mechanisms driving the mouse-tracking paths and the implications for the measurement of deep-parameters like tTaste and tHealth. For example, based on our continued work in this area, we now view the link between the state of an integrator in a DDM and the mouse-tracking movements in our original model as too simplistic, and are currently building improved computational models of the mouse-tracking task and its implications for experimental design and measurement. We are hopeful that this line of work will result in substantial improvements in the experimenter’s ability to measure the latency of various types of information during decision making.

Unfortunately, however, the key criticisms C1-C3 made in the current piece are flawed, as we explain in detail below.

### Problems with C1

Criticism C1 regarding the choice confound is incorrect. In fact, adding the proposed choice direction regressor eliminates the ability to correctly measure differences in the latencies of tTaste and tHealh even when they exist <u>by assumption</u> in simulated data.

The intuition for the problem is simple to see by considering a simpler problem. Imagine that you have binary choice data that <u>by assumption</u> is generated by a standard logit process, so that the probability of choosing left is a logit function of the latent value, *v*, of the item. If you run a logit regression of choices on *v* you will get an unbiased estimate of the true impact of value on choices. However, if, following the authors’ proposal, you run a regression of choice on value and an additional variable that is highly correlated with actual choice, you would falsely conclude that the estimated impact of the latent value is negligible. This same problem is present in the authors’ C1, applied to the mouse-tracking regressions one time point at a time.

To illustrate the problem even more starkly, we have simulated a mouse-tracking dataset where <u>by assumption</u> the latency of tTaste is faster than the latency of tHealth. This shows that whereas reasonable estimates of tTaste and tHealth are obtained with the method from Sullivan et al. (2015), introducing the choice regressor proposed in C1 instead produces incorrect estimates of these latencies.

The details of the simulation can be seen in the published open access code (CodeforC1a.R and CodeforC1b.R). The idea of the model is simple and illustrated in Figure 1. Subjects move their cursor straight up at constant velocity until they decide which option to choose. Once a decision is made the cursor moves in a straight line to the chosen option. The duration of the upward movement is determined by a Drift Diffusion Model (DDM) which weights the relative value of the health and taste attributes and allows for different latencies in the computation of these variables.

**Figure 1.**
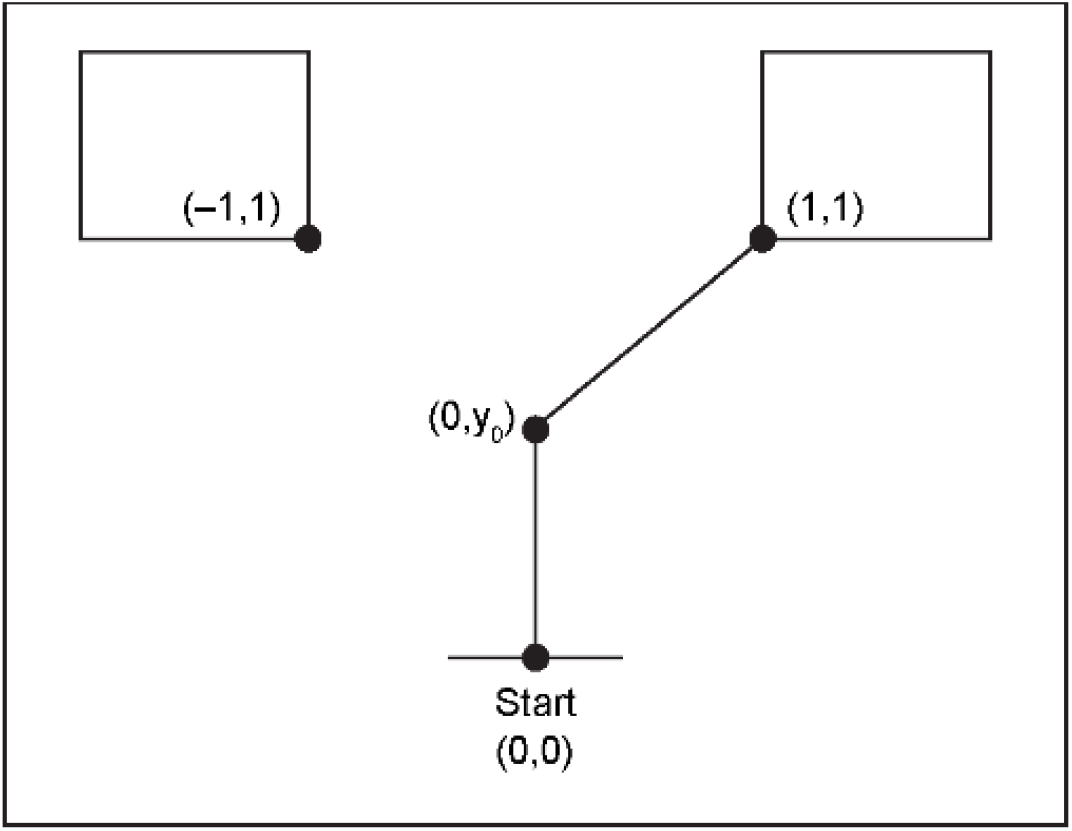
Illustration of the simulated model for C1.

We simulated data for an individual using this model. This resulted in simulated mouse trajectory paths, which we then used to carry out mouse-tracking path regressions as in Sullivan et al. (2015) with and without controlling for choice. In Figure 2, panels A and B show the results in a situation when <u>by assumption</u> the latency for taste is faster than for health (tTaste = 250 ms, tHealth = 500 ms), and panels C and D show the case when <u>by assumption</u> the latency for taste is equal to that for health (tTaste = tHealth = 500 ms). The estimated paths of the influence of taste are depicted in red, those for health in blue, and those for the choice regressor in black. The figures show that the method in Sullivan et al. (2015) generates reasonable estimates of tTaste and tHealth, but that adding the choice regressor strongly biases the coefficients to zero and eliminates the ability of the method to measure the latencies.

**Figure 2.**
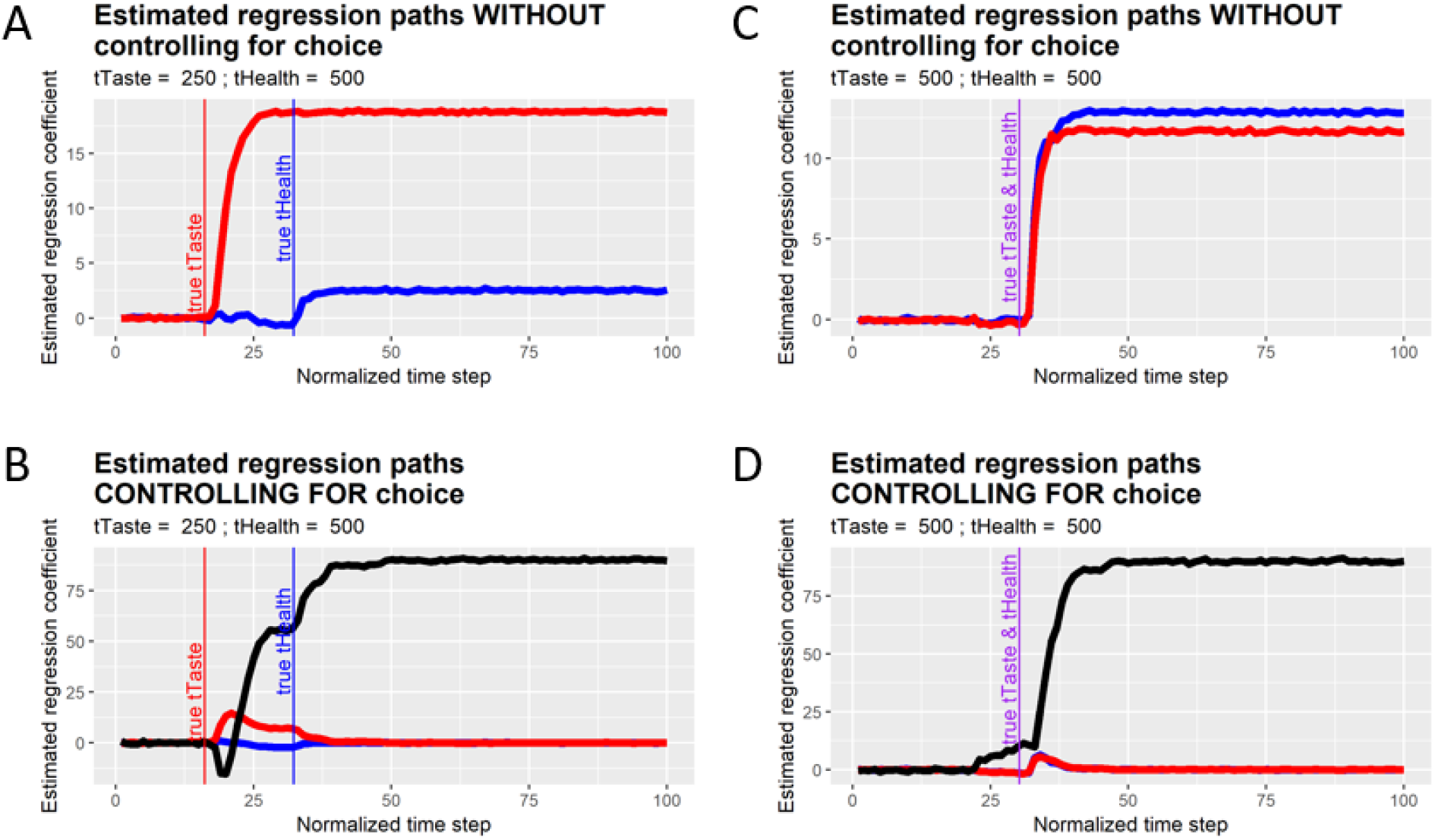
Estimated regression paths using simulated data.

### Problems with C2

Criticism C2, regarding the relationship of the curvature in the filler task and the estimates of tTaste and tHealth, also contains a fundamental flaw.

Zhang et al. (2018)’s criticism is based on the <u>assumption</u> that there should be no relationship between taste and health latencies and the maximum deviation curvature measure. For example, in p.6 they state that “as the filler task does not require cognitive processing of healthiness and tastiness, one would not expect any correlations between estimated processing speed and size of the curvature in these filler trials”, and in (p. 16) they state that “if the processing speeds in Sullivan et al. (2015)’s method were derived from the genuine influences of attributes, they should not correlate with personal mean MD (their curvature measure) in the filer trials”

But as the following example shows, one would expect such a correlation even in very simple models, and its existence need not confound the estimates of tTaste and tHealth generated by the method in Sullivan et al. (2015).

The details of the example are presented in the open access code (CodeforC2a.R and CodeforC2b.R). But the intuition behind the example is simple (see Figure 3 for illustration). We simulate data for 50 different subjects that make mouse-tracking movements using the model above and that differ <u>only</u> on the velocity at which they move the cursor upwards during the initial phase. We also simulate data from these same subjects using a filler task, with the assumption that the same velocity parameter is at work for each individual in both tasks. We measure the curvature of the paths using the same maximum deviation (MD) measure utilized by the authors.

**Figure 3.**
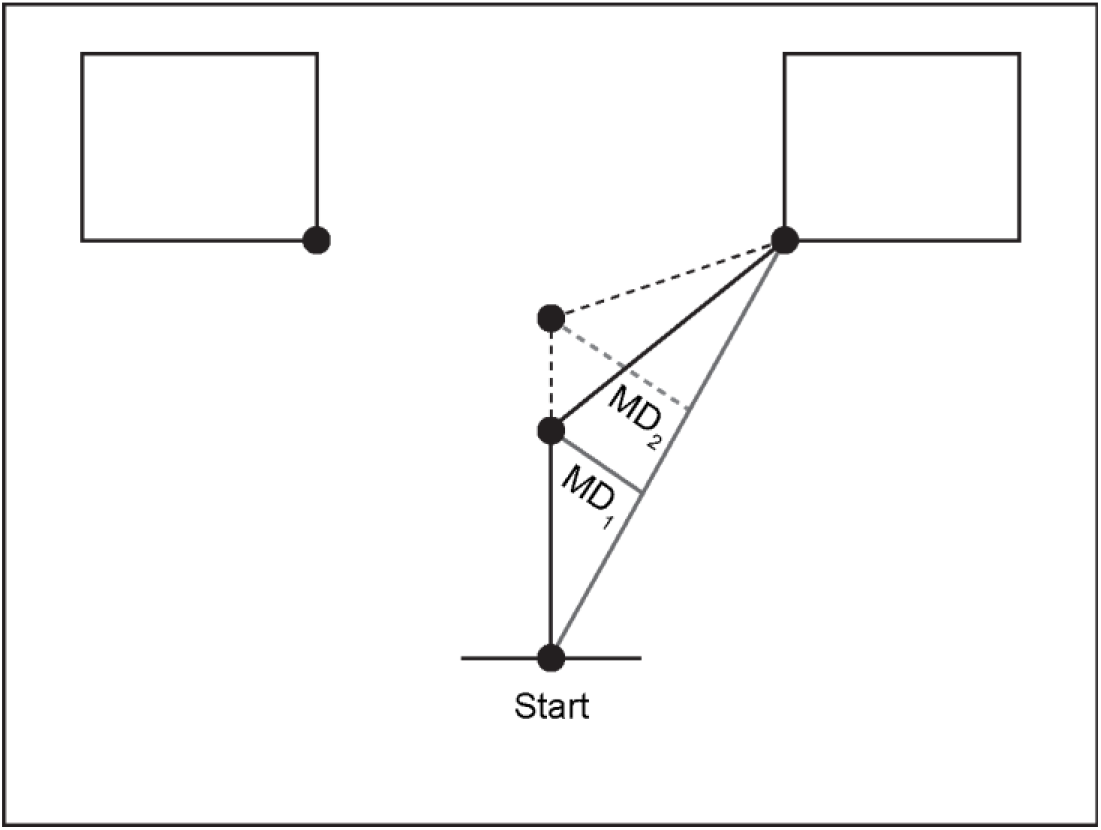
Illustration of the simulated model for C2.

As before, we do two sets of simulations and present the results in the next figure. Figure 4 panels A and C show the results in a situation when <u>by assumption</u> the latency for taste is less than for health (tTaste = 250, tHealth = 500). Panels D and F show when <u>by assumption</u> the latency for taste is equal to that for taste (tTaste = tHealth = 500).

The results show that the assumptions behind C2 are incorrect. Figure 4 panels A and D show a high degree of correlation between the MD measure in both tasks. Panels B, C, E and F show a correlation between the MD measure and the estimated latency times in a model, as one would expect since velocity affects the vertical location of the path kink and the MD measure is a linear transformation of the kink location. More importantly, a comparison of panels B and C vs. E and F shows that the differences in velocities do not affect the relative estimates of tTaste and tHealth: tTaste is estimated to be about half of tHealth when <u>by assumption</u> tTaste = 250 and tHealth = 500 and is estimated to be approximately equal to tHealth when <u>by assumption</u> tTaste = tHealth = 500.

**Figure 4.**
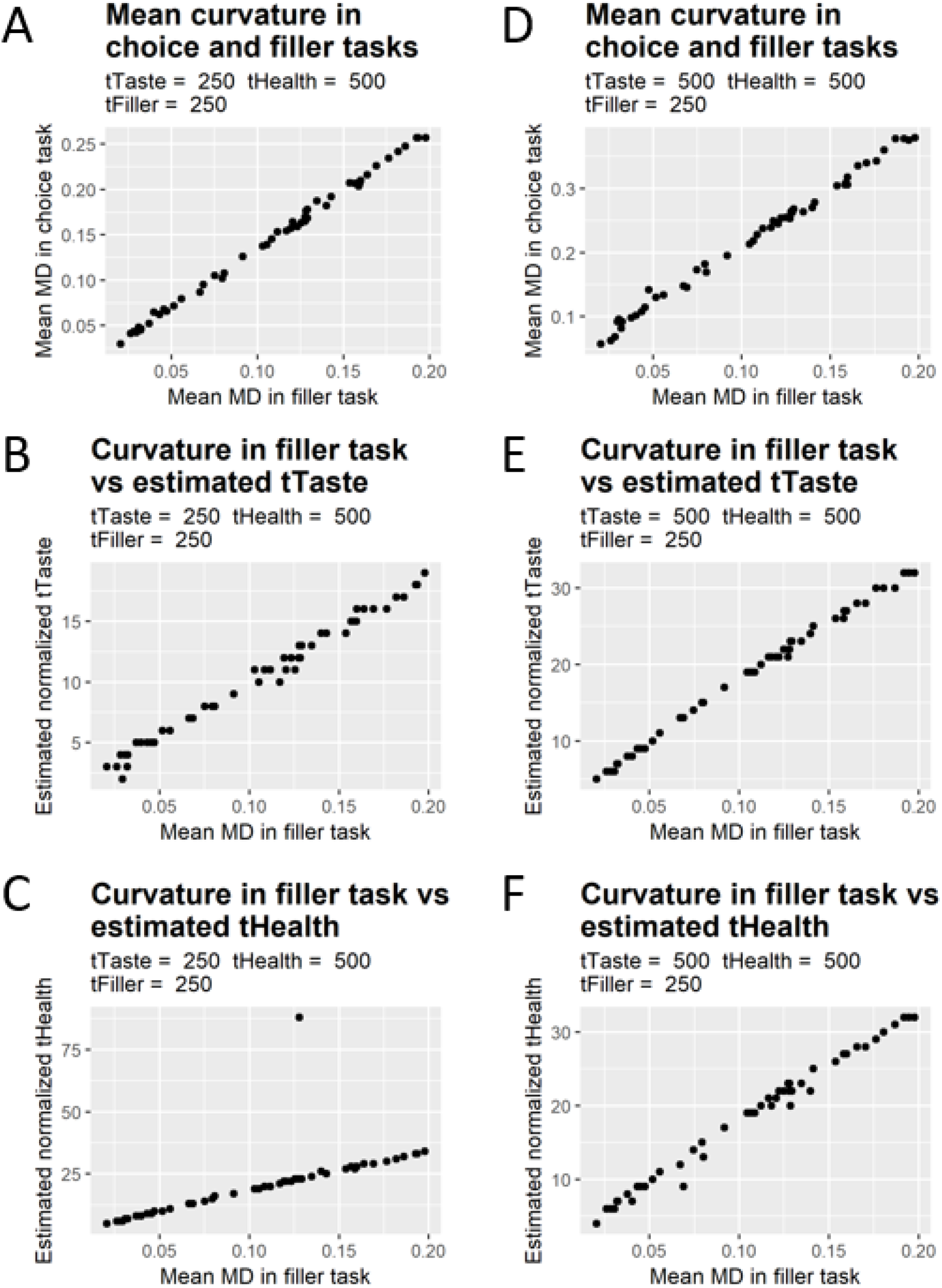
Maximum deviation (MD) correlations with latency parameters.

### Problems with C3

We also have concerns with criticism C3, regarding the alleged lack of methodological robustness of one of the statistical methods used in Sullivan et al. (2015).

The authors do not provide any direct criticism of the conceptual validity of the method. Instead, the essence of their criticism is that the method does not seem to work as intended in their own dataset.

The problem with this argument is that the method can fail to work as intended not because of an inherent lack of robustness, but due to problems or inconsistencies in the experimental data to which it is applied.

Unfortunately, this appears to be the case with the study that the authors used to make C3, which has flaws in experimental design that can create problems in measuring tTaste and tHealth. In particular, the authors only used 10 food stimuli with highly negatively correlated health and taste ratings. This gives rise to a multi-collinearity problem that can impede the correct estimation of the mouse-tracking path regressions. In contrast, in Sullivan et al. (2015) the health and taste ratings are decorrelated by design to avoid this problem.

To illustrate this more precisely, we carried out another simulation using the same model as above, but with health and taste negatively correlated as in the authors’ experimental design (details are presented in the open access code CodeforC3.R). As shown in Figure 5, this results in “negative estimates” for the influence of health during a sizable part of the decision period, even though <u>by assumption</u> health has no effect on the mouse-tracking paths during this period.

**Figure 5.**
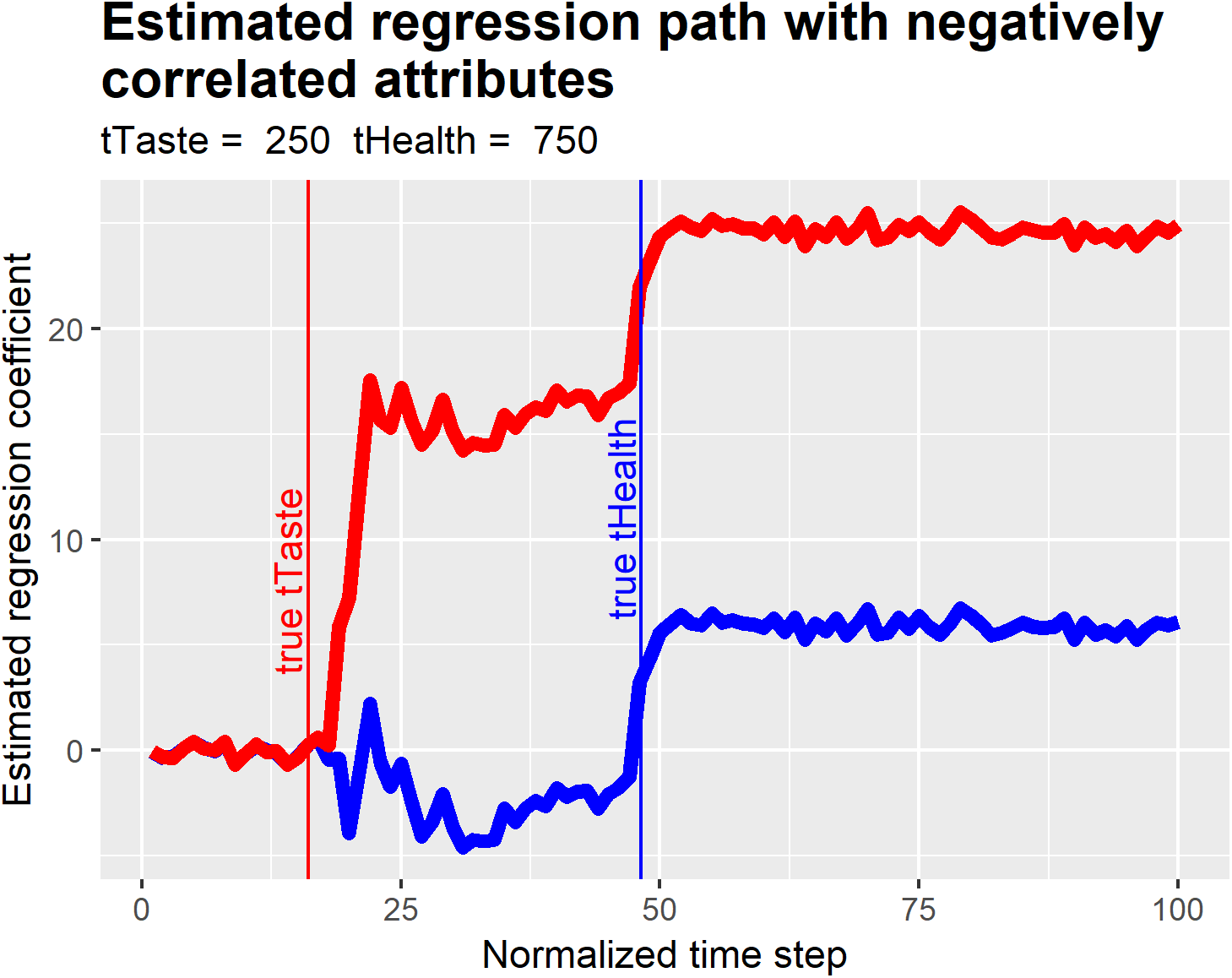
Estimated regression paths for taste and health when attributes are negatively correlated

In addition, unlike Sullivan et al. (2015), which estimates the paths for health and taste using a single regression, an examination of the authors’ code suggests that they have run independent path regressions for taste and health, which further exacerbates these estimation problems. Together, these problems cast serious doubt on the reliability of C3.

### Other problems

Although the above comments concern the authors’ criticisms of the mouse-tracking methodology in Sullivan et al. (2015), it is also important to note that our results for the relative latency of tTaste and tHealth have been replicated in other studies using estimates that *depend only on choice and RT*, without using the mouse-tracking path data.

Consider, in particular, the study by Maier et al. (2018) and Sullivan et al. (2018). Both papers estimate tTaste and tHealth using <u>only</u> choice and RT data, as shown in the below figure from Maier et al. (2018). Consistent with Sullivan et al. (2015), they find a shorter latency for Taste than for Health information, which is replicated in other datasets with different participants and task demands.

Together with the simulation data described above, these results provide strong evidence against the claim that the findings of Sullivan et al. (2015) are the result of a “motor-confound”, as argued by Zhang et al. 2018.

**Figure 2.**
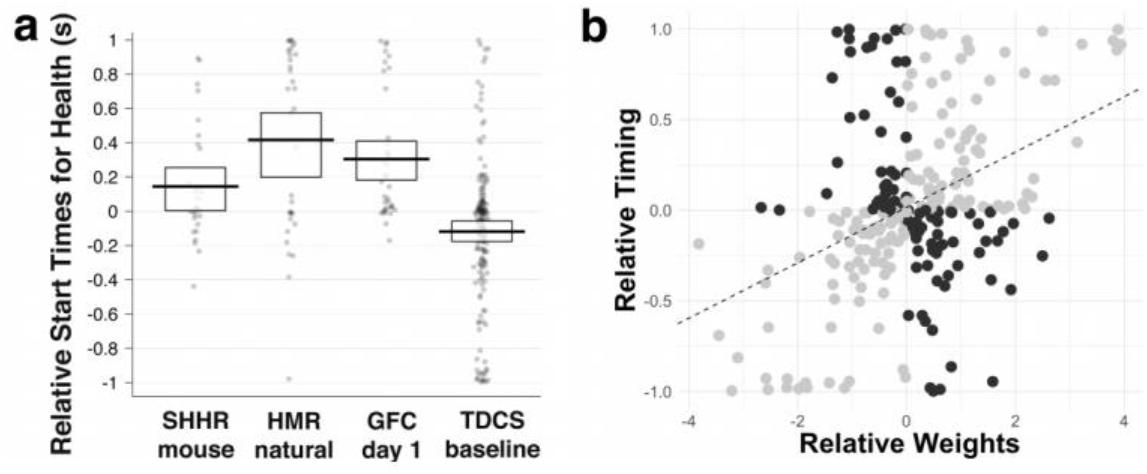
Panel **(a)** shows the relative start times in seconds for healthiness compared to tastiness for all participants In each study. Positive values indicate that tastiness is considered before healthiness and negative values that healthiness Is considered before tastiness. In each column every dot Is a separate participant. The thick black horizontal bars represent w¡thin-study means and the rectangular bands indicate the 95% highest density intervals (HDIs). Dataset abbreviations: SHHR = data from the computer-mouse response trials In Sullivan et al 2015; HMR = data from the natural choice condition In Hare et al 2011; GFC = newly collected data from the first session/day of an experiment combining gambles and food choices; TDCS = newly collected data from the pre-stimulation baseline choices in our tDCS experiment. The scatterplot in **(b)** plots each participants’ relative timing data against attribute weights, separated by whether the relative weighting of tastiness and healthiness and their relative timing are aligned (gray circles) or whether there is misalignment between weighting and timing (i.e., the highest weighted is not the fastest; black circles). The highest weighted and fastest-to-be-considered attributes were misaligned in 38% of participants.

